## Supplementary File 1 for "A Bidirectional Network for Appetite Control in Larval Zebrafish"

### <u>Conceptual Circuit Model</u>

The core issue that arose during the review of the manuscript, and the apparent paradox that manifests in the observant reader, is that cH activity correlates both with Hunger and Satiety - depending on the presence or absence of food. This conundrum is also reflected in the reviewers' comments.

At a more basic level, there was also some concern about usage of words such as hunger and satiety in larval zebrafish, which we believe is largely a semantic issue that is best addressed by providing an operational definition of these terms:

<u>Hunger</u> is an internal state defined by three conditions:
1) The state of being in a nutrient/energy deficit - which is usually the consequence of food deprivation
2) A state that may promote food-seeking behavior as well as increased food intake
3) A state that is reflected by - and correlates with - a pattern of modulatory neuronal activity

<u>Voracious feeding</u> is a sub-state of hunger in which food is present and being ingested, but nutrient/energy deficit is still high.

<u>Satiety</u> is considered the opposite of hunger and is defined accordingly:
1) A state of having sufficient levels of nutrient/energy or levels that rest above a homeostatic baseline
2) A state that manifests behaviorally in the slower/lower consumption of food relative to the state of hunger, due to indifference to food or its active avoidance
3) A state that is reflected by an internal modulatory neuronal state that may be antagonistic and opposite to that from hunger

In order to clarify these questions related to our manuscript we give in the following a detailed description of our conceptual model. We describe how this model incorporates the observed activity patterns in all three nuclei ( cH, mLH and lLH) and we discuss the role that we propose all three may play in releasing hunting and feeding behavior. **We emphasize that this is a working model, and though we have presented partial evidence in support of some aspects, additional studies will be required to conclusively prove and/or refine our hypotheses.**

To recap, we observe that cH activity is at baseline when the animal is satiated (well-fed). Activity then increases during food-deprivation, and drops to very low levels once food is presented. During feeding, and with the resulting return to satiety, the activity rises back to baseline levels.

**Figure 1: Summary of cH and LH activity over Hunger and Satiety**

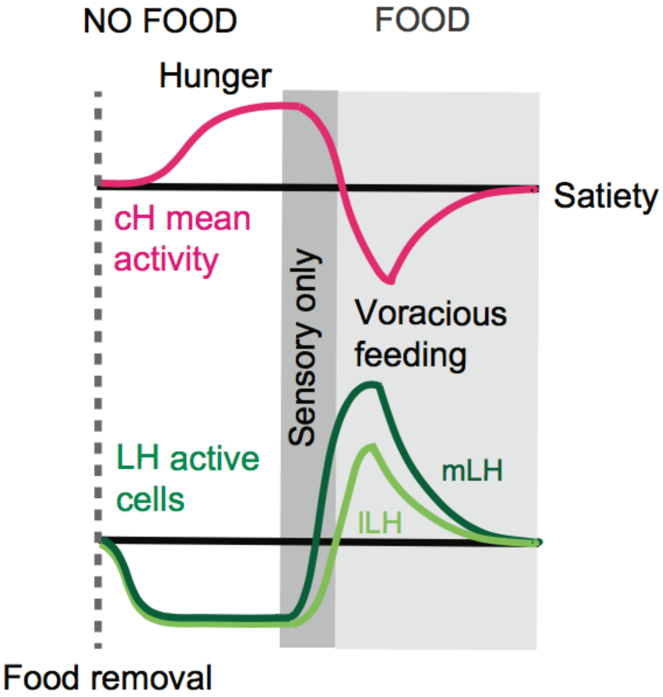


The general framework of our conceptual circuit model is based on two core assumptions. These are:

1) Similar to mammals and adult fish, **the LH drives food consumption** (Demski and Knigge, 1971; Jennings et al., 2015; Roberts and Savage, 1978)**.** Thus, it makes sense that LH activity *in the presence of food/ingestive cues* is at medium levels during satiety and higher with increased food-deprivation time.
2) That the cH and the lLH/mLH have **mutually inhibitory connectivity**, a hypothesis which is backed up by the strong and consistent anti-correlated activity patterns observed in these nuclei. **Further, we now provide evidence demonstrating an inhibitory effect of the cH on lLH activity.**

We will also be making other assumptions which are explicitly outlined below.

Next, we address the seemingly-complex role of the cH, which is complicated by the fact that it has opposite activity patterns depending on whether food is absent or whether it has been detected/ingested. In our manuscript we put forth a number of non-mutually-exclusive hypotheses for the roles of the cH in each of these stages:

1) **The cH encodes an aversive, negative valence state,** which should induce, when activated in the absence of food, a negative association with the current condition and a drive to change this condition and explore alternatives. Activating cH in the presence of food should reduce food intake, as observed with other stressors (De Marco et al., 2014), and optimize the animal's behavior to remove itself from the negative context. For

example in mammals, AgRP neurons, which are activated during hunger, have been shown to encode a similar aversive state (Betley et al., 2015).

2) **The cH drives exploration (food search) as *opposed to* exploitation (food consumption),** which could be reflected by enhanced locomotion, increase in search area, enhanced visual acuity or sensitivity to prey-like objects or other subtle changes that might not manifest significantly in spontaneous swimming behavior in zebrafish larva. When activated in the absence of food it might drive enhanced exploration and higher sensitivity to food-like cues. For example in mammals, AgRP neurons promote food seeking and exploratory behaviors (Dietrich et al., 2015; Krashes et al., 2011). In contrast, activating the cH in the presence of food, while possibly lowering the threshold of initiating food-seeking behavior (i.e. exploration), might ultimately lead to a reduction of food intake and consummation (i.e. exploitation).

3) **The cH induces sensitization/priming of the LH circuit** such that it is more responsive to food and ingestive cues. If accurate, when activated in the absence of food it should drive enhanced feeding after food is presented. **This is what we have observed and reported in our manuscript (Figure 6).** If the cH is activated in the presence of food, as long as the priming effect is weaker than the acute cH-mediated inhibition of the LH, driving the cH should simply reduce LH activity and thus also reduce consummatory behavior. Otherwise, if the priming effect is strong, consummatory behavior might be increased. **Our experiments activating the cH in the presence of food in both fed and food-deprived fish suggest that any effect of priming is weaker than the acute inhibitory effect of the cH on LH activity.**

In the current manuscript, we present partial evidence in support of all three hypotheses. We want to make clear that a conclusive verification of the complete conceptual circuit model (see below) is beyond the scope of this manuscript and will require future studies.

**<u>A putative circuit diagram</u>**

We postulate that the cH receives excitatory input from an undefined source (possibly even directly sensing nutrients/hormones from ventricular cerebrospinal fluid) that generally codes for caloric deficit. We will call this the hunger encoding circuit (HEC). In addition we postulate that the cH receives excitatory input from a source analogous to the HEC that generally codes for satiety. We'll call this unknown circuit the "satiety encoding circuit" or "SEC". It is possible that different neurons within the cH encode each of these cues, but both need to be represented for cH activity to converge stably at "medium" levels during satiety (where HEC = 0) without drift.

As described before, we propose that the cH also receives inhibitory input from the LH, which is gated by food and ingestive cues, as well as by satiety cues. Thus, in the absence of food, the LH is inactive and the HEC activates the cH, whereas in the presence of food, the cH is initially strongly inhibited by the LH, and then, as the HEC is shut down and the SEC activated, returns to a baseline firing rate.

Finally, on top of this basic circuitry we propose a latent modulatory connection between the cH and LH which primes the sensitivity of the LH during hunger such that it becomes more active once food and ingestive cues are presented. Other yet-to-be-discovered HEC components may also be involved in such priming. Details of the model are outlined below.

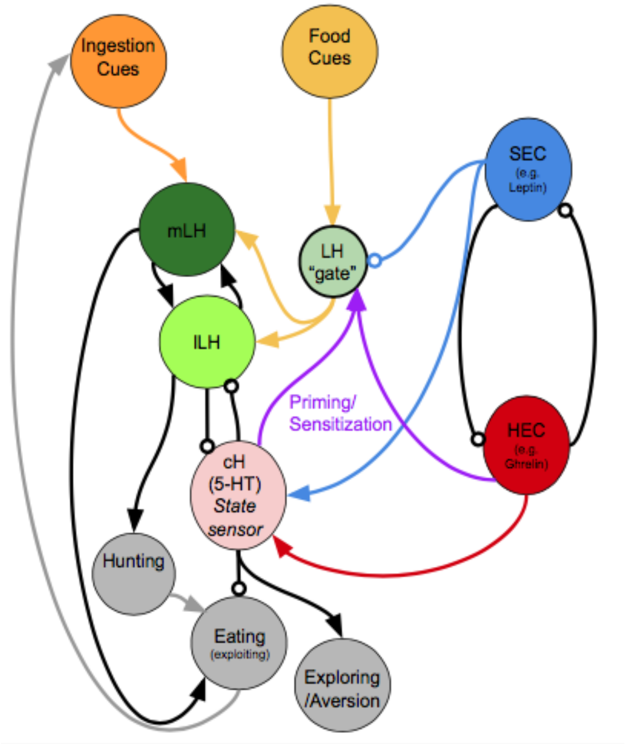


**Figure 2: A putative circuit diagram**
HEC = Hunger Encoding Circuit, SEC = Satiety Encoding Circuit, which should have anti-correlated activities and report the animal's energy/caloric status. The cH represents both hunger and satiety state and primes the LH during hunger. It may drive other behaviors such as exploration or aversive behavior, but also suppresses feeding. Other HEC components may also be involved in LH priming/sensitization. We propose mutual inhibition between the cH and LH, though we have only demonstrated unidirectional inhibition (cH on lLH) thus far. The mLH, normally responsive to food cues, may promote hunting, though not necessarily coupled with ingestion, whereas the lLH, which is more responsive to ingestive cues, should enhance further ingestion (i.e. eating). The LH "gate" is a conceptual representation of how its sensitivity to food cues could be modulated by other signals (i.e. reduced by the SEC and enhanced by cH-mediated priming). It does not necessarily represent a physical neuronal population.

Below we discuss in detail the prediction that this model generates for targeted silencing and activation of cH in *food-deprived* as well as *fed* fish in both, the *presence* as well as the *absence* of food. Note that there are 8 possible optogenetic experiments to be conducted with the cH as a target, and they are comprised of three pairs of mutually exclusive conditions: activation vs silencing, food-deprived fish vs fed fish, presence of food vs absence of food. Food intake is currently the clearest behavioral readout, and thus also the focus of this manuscript, though we will continue to pursue experiments in support of other predictions (e.g. exploratory behavior, aversive conditioning).

An additional 16 experiments are possible if the mLH and lLH are also considered possible targets for specific optogenetic perturbation. However, such experiments are currently difficult to do since specific transgenic lines for these regions are not available.

**<u>The serotonergic neurons of the caudal hypothalamus (cH):</u>**

We propose that the cH is driven by (i.e. receives excitatory input from) two regulatory centers: the hunger encoding circuit (HEC), and the satiety encoding circuit (SEC), whose existence we postulate, but which we currently cannot back up with supporting experimental data. Thus, the cH can be strongly activated by hunger cues, but also returned to baseline levels by satiety cues after having been strongly suppressed by the LH.

As described above, we postulate that the cH may play multiple roles in regulating feeding behavior. One of these roles may be to "prime" the LH circuit such that it becomes more active once food is presented. This postulate is supported by the observation that the amount of food-cue-induced activity in LH increases with food deprivation time and hence with increased integrated cH activity during food deprivation.

Once food is presented, we posit that the LH, driven by food and ingestive cues, shuts down the cH via inhibitory circuitry. It does so more strongly in food-deprived vs fed fish, as it has been primed by prior cH/HEC activation, and also the SEC is not activated (i.e. the putative "gate" is wide open). This is supported by evidence that LH activity is much higher after a longer-period of food-deprivation (e.g. Manuscript Figure 2).

In the presence of food, recovering cH activity levels during feeding lead to a progressive reduction in LH levels and thus reduce food intake.

<u>Predictions:</u>

*Activation* of the cH in the *presence of food* should reduce ingestion and feeding rates. **This is consistent with our current optogenetic results in food-deprived fish.** In the case of satiated fish (an experiment we did not previously attempt), we predicted that food intake should still be reduced, but counteracted by the effect of priming. We now present results consistent with this hypothesis-- food intake trends lower in continuously fed fish, though not significantly.

Activation of the cH in the *absence of food* may lead to 1) avoidance and/or aversive conditioning towards the location at which the cH is being stimulated 2) increased exploratory behavior; e.g. increase of swim frequency, subtle convergence of eyes, or other subtle changes that might not manifest significantly in spontaneous swimming behavior in zebrafish larva.

Activation of the cH in the *absence of food* may also sensitize the LH circuit to future food cues and enhance food intake. **This is consistent with our optogenetic results.**

Ablation of the cH should lead to a disinhibition (i.e. a release) of LH activity and thus hunting and ingestion. Though any "priming" effect would now be reduced, we posit that the acute disinhibition of LH activity would drive up food intake regardless of whether it has been previously "primed" by the cH. That is, animals should still eat more in the presence of food. **This is consistent with our ablation results.**

In the absence of food we expect that fish with an ablated cH would display a reduction in exploratory behavior which might manifest in a decrease in swim frequency, divergence of eyes etc; again, this might be hard to pick up because these expected changes are subtle and might not present yet at larval stages.

**<u>The neurons of the lateral part of the lateral hypothalamus (lLH):</u>**

We observe that the lLH shows clear anti-correlated activity with the cH. Its activity is at baseline during satiation, activity decreases during food deprivation, switches to high levels when the animal starts feeding, and slowly drops back to baseline level with the return of satiety.

We postulate and now demonstrate using optogenetics that the lLH likely receives inhibitory input from the cH, and that it is also strongly excited by consummatory internal cues, e.g. food being ingested. An example of such an input is the anterior branch of the esophageal nerve (En2) in Aplysia which is both necessary and sufficient for effective reinforcement of biting behavior (Brembs et al., 2002) and which is known to convey information about the presence of food during ingestive behavior. Based on work from Muto et al (2017), LH activity (they did not distinguish between the lobes) increases slightly after food detection, but even more strongly immediately after ingestion (Muto et al., 2017). We further propose that activation of lLH only occurs if both inputs are activated together (cH inhibition and food ingestive cues) and that its activity drives ingestive behavior (capture swims, biting and swallowing).

<u>Predictions:</u>

Since no specific transgenic lines exist, clean perturbation experiments are not possible; but predictions, of course can be made.

*Activation* of the lLH should lead to voracious feeding in the presence of food.

*Silencing* of the nucleus should shut down consummatory behavior.

**<u>The neurons of the medial part of the lateral hypothalamus (mLH):</u>**

We observe that the mLH basically shows the same activity patterns as the lLH. The only difference is that the switch from depression to activation occurs already with the presentation of food cues, though there is also a further enhancement post-ingestion. This is concluded from: 1) our calcium imaging results with live paramecia (which fish are unable to consume); 2) the fact that exposure to artemia as food cues, which fish can hunt but cannot swallow (they are too big) drives activity in mLH but not in lLH.

We also have evidence (not included in the manuscript) that the mLH is responsive to paramecia odor. In addition, evidence of LH activation by vision was also presented by Muto et al (2017). We thus postulate that the mLH receives strong excitatory input from the sensory modalities that detect food (vision, olfaction etc), and that it is inhibited by the cH.

We further postulate that the mLH drives the transition from exploratory to hunting behavior, i.e. it drives the initiation of hunting sequences that start with re-orienting turns (j-turns) and eye-convergence and are followed by pursuits of prey.

<u>Predictions:</u>

Similar constraints about the implementation apply, but predictions can be made.

*Activation* should lead to an increased probability of the initiation of hunts vs exploratory swims.

*Silencing* should do the reverse, namely induce a relative reduction in the release of such hunting sequences.

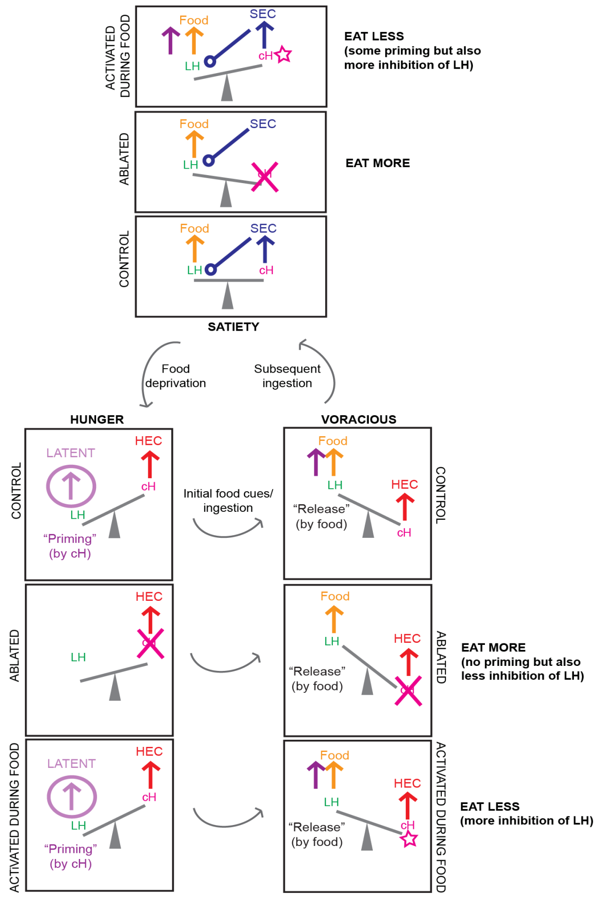


**Figure 3: Another schematic summarizing our circuit model, including predictions for cH ablation and activation results.** Here, the proposed mutual inhibition between cH and LH activity is represented by a "see-saw", and relevant inputs/outputs during each phase are represented by "forces". We represent our predictions for what happens when the cH is ablated, as well as when it is activated in the presence of food, for both satiated and hungry fish. We assume that the effects of priming are weaker than the effects of acute mutual inhibition. Color codes are consistent with Figure 2. Note that activation of the cH *prior* to feeding is not depicted in this diagram: in this scenario we predict that this would cause priming of the LH that increases subsequent feeding, the effect of which may be more obvious in satiated fish where cH starts off lower (consistent with our optogenetics results).
